# Mouse hue and wavelength-specific luminance contrast sensitivity are non-uniform across visual space

**DOI:** 10.1101/174631

**Authors:** Daniel J. Denman, Jennifer A. Luviano, Douglas R. Ollerenshaw, Sissy Cross, Derric Williams, Michael A. Buice, Shawn R. Olsen, R. Clay Reid

## Abstract

Mammalian visual behaviors, as well as responses in the neural systems thought to underlie these behaviors, are driven by luminance and hue contrast. With tools for measuring activity in cell-type specific populations in the mouse during visual behavior gaining traction, it is important to define the extent of luminance and hue information that is behaviorally-accessible to the mouse. A non-uniform distribution of cone opsins in the mouse potentially complicates both luminance and hue sensitivity: opposing gradients of short (UV-shifted) and middle (blue/green) cone opsins suggest that hue discrimination and wavelength-specific luminance contrast sensitivity may differ depending on retinotopic location. Here we ask if, and how well, mice can discriminate color and wavelength-specific luminance across visuotopic space. We found that mice were able to discriminate hue, and were able to do so more broadly across visuotopic space than expected from the cone-opsin distribution. We also found wavelength-band specific differences in luminance sensitivity.

## Introduction

The mouse visual system is increasingly ^1,2^ being used as a model system for studying both cortical sensory processing ^3–7^ and behavior ^8–12^. While most physiological work has used achromatic stimuli^3,13^, mice, like most other mammals, display physiological color-opponent signals in the retina ^14–18^, through LGN ^19^ and possibly V1 ^20^. The mouse retina displays asymmetric and mixed expression of its two opsins along the dorsal-ventral axis of the retina, creating opposing gradients of short and middle opsins ^21,22^ and resulting in gradients of wavelength-band specific responses ^16,19,23^. Therefore the substrate for cone-driven color-opponent signals, and any hue sensitivity, exists only in the overlapping “opsin transition zone” ^15,19^. However, short and middle opsin responses broadly overlap in V1 and higher visual areas ^23^ and rod-cone antagonism can also create color opponency in some mouse retinal ganglion cells ^17^, presenting the possibility that behaviorally-relevant color information could be extracted more broadly across retinotopic space.

Whether mice can use hue information to guide visual behavior is an open question. There is some evidence for hue discrimination ^24^, but it remains unclear how this depends on overall luminance, luminance contrast, or retinotopic position. Further, it is not known if the gradients in opsin distribution lead to variations in behavioral luminance sensitivity across space. Such non-uniformity would impact studies of visuotopically extended V1 populations, such as studies of population sparsity ^25^, population correlations ^26^ and other notions of population coding ^10,27^.

Here we use a simple behavior, change detection, to determine where in visual space mice can discriminate changes in hue and luminance at ethologically-relevant (i.e, mesopic) luminance levels. By measuring detectability of luminance and color changes separately across elevation (spanning ∼75°), we are able to generate an estimate of wavelength-specific contrast sensitivity across visual space. Mice were able to discriminate hue, but only at elevations above the horizon. We find both wavelength-specific luminance and hue contrast sensitivity to be dependent on retinotopic location, but that these differences in sensitivity were less dramatic than expected from the cone opsin distribution, suggesting behavioral access to differential activation of rods and cones.

## Results

### Behavioral Task

To examine the psychophysical and physiological basis of mouse color vision, we first trained mice in a go/no-go change detection task^8^ in an immersive visual stimulation environment customized for delivering stimuli in the spectral bands of the mouse short and middle wavelength opsins (Fig 1A; *Materials and Methods*). We use the system here to deliver a video stimulus driven by a green and ultraviolet LED projector; for each point on the stimulus the green and ultraviolet intensity could be independently modulated. Total luminance was in the mesopic range, over which mice are both behaviorally active ^28^ and color opponent signals have been demonstrated in the retina ^15,17^. Under this paradigm, mice indicate that they have perceived a change in the stimulus by licking a reward spout within 1 second of the change (Figure 1B); subsequent licks allow reward consumption (Figure1C).

**Figure 1.**
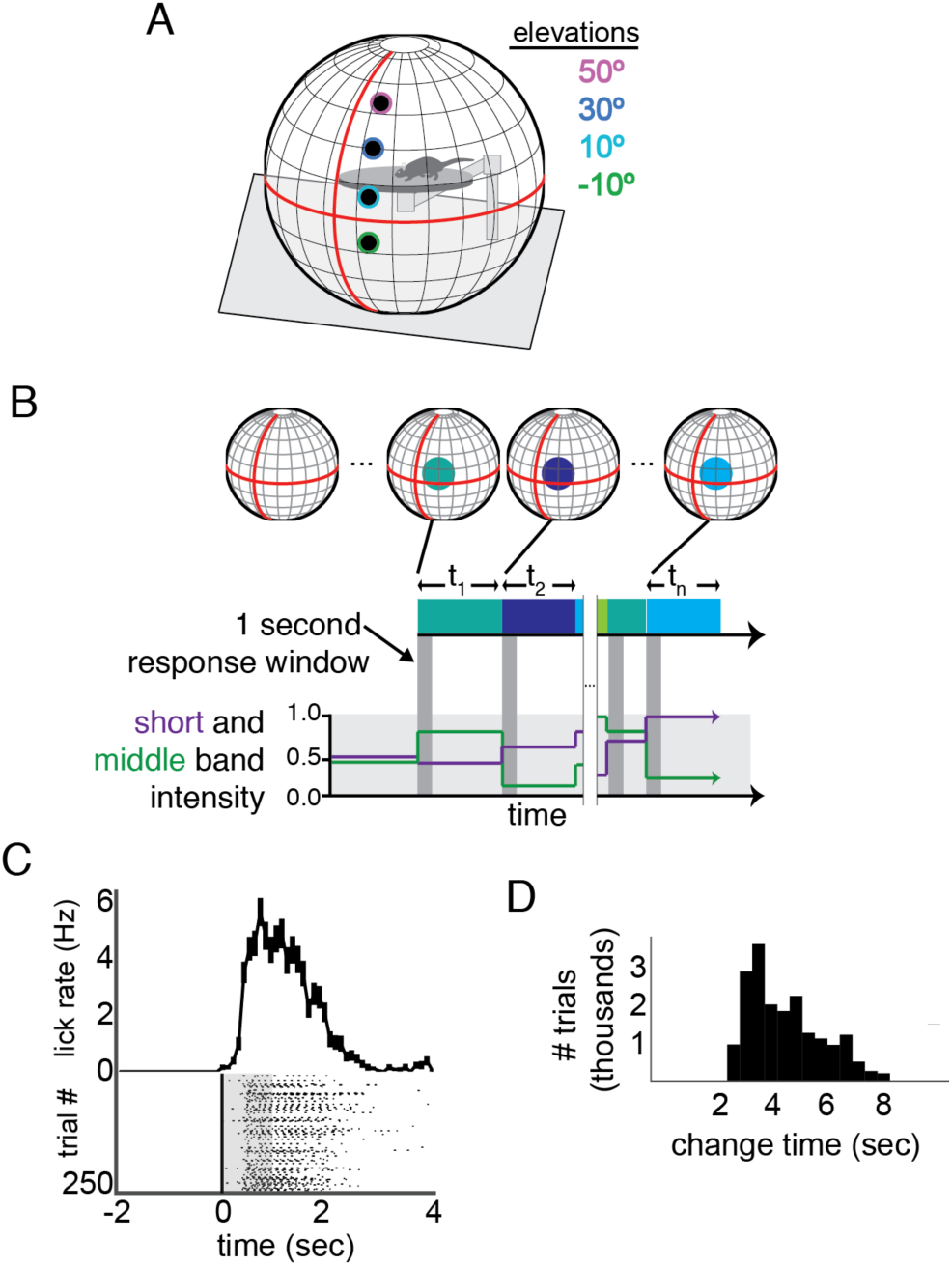
Change detection task in an immersive visual stimulation environment capable of delivering short-and middle-wavelength band stimulation. **A**, The visual stimulation environment, with the positions and size of stimuli shown; the colored edges were not a part of the presented stimulus, but indicate the color scheme used to denote elevation throughout the other figures. **B,** A schematic of the task. The background was set mean intensity for each wavelength band. At variable times, t_n_, the intensity of short and middle wavelength bands within a 15° diameter circle changed. If the mouse licked within 1 second of this change (indicate by the dark grey boxes), a reward was delivered. Schematic short and middle band intensities are shown on the lower plot and corresponding stimulus changes for each epoch are schematized in blocks and projected as circles onto a sphere. The schematized circles are larger than actual stimuli, for clarity. **C,** Example performance in a single session across 250 trials. Each lick is shown relative to stimulus change; the response window is overlaid in grey. A histogram of lick times is shown above. Error bars are S.E.M. **D,** The distribution of change times (t in panel **B**) across the trials used for analysis of change detection performance. The distribution follows the log sampling distribution, enforcing roughly equal probability of a change occurring as the mouse continues to wait.

Following pre-training on the change detection task (see *Materials and Methods*), we switched to change detection sessions in which the ultraviolet and green intensity, centered on the mouse short and middle wavelength bands, respectively, were varied independently on each trial. Each trial contained a change in intensity for a 15° test circle on a mean luminance background at one of four elevations: -10°, 10°, 30°, and in some cases 50° (relative to both the horizon and the placement of the rotating mouse platform). We varied position only along elevation because both rods and cones are relatively uniform across the azimuthal axis of the retina^29^. Eye position did not change with stimulus location (Figure 1 – figure supplement 1). To achieve sufficient trials to cover this stimulus space, we presented a total of 127659 trials (n=4/5 total mice trained, 284 sessions). To control for motivation, we calculated a running average of the reward rate and selected trials where this reward rate remained above 4 rewards per minute; only these engaged trials (44%, 56112/127659) were used for analysis.

### Short and middle wavelength band specific contrast sensitivity

We first examined our results to estimate the relative luminance contrast sensitivities to short and middle-wavelength band stimulation across the visual field. Although the green and ultraviolet projector LEDs nearly isolate responses of the middle and short wavelength sensitive opsins^30^, they do not necessarily isolate responses of individual cones, most of which express a combination of the two opsins. Nor is it necessarily a measurement of the relative weight of the cone opsins themselves, as rods also may contribute to this light sensitivity at these luminance levels. Rather, we present a measure of the relative perceptual weight to stimuli of the middle and short wavelength bands covered by our stimulus LEDs (Figure 2 – figure supplement 1), as combined through both cone opsins and rods.

Total luminance change detection saturated by ∼30% at all elevations (Fig 2A) and the half-saturation threshold (hereafter referred to as “threshold”) < 12% for each elevation (Fig 2B). The highest sensitivity was in the upper visual field, at 5.5% threshold, a threshold that is consistent with previous reports for a 15° (∼0.07 cyc/°) stimulus^8,31–33^. Sensitivity to increments in contrast was similar for all elevations (10.5-12.1%). Consistent with previous physiological measurements in V1 of mouse ^20^ and other species^34,35^, sensitivity to decrements in contrast was higher than sensitivity to increments (5.4% - 10.4%). Notably, this was most pronounced in the upper visual field (difference: 5%) than the lower visual field (difference: 1.3%).

**Figure 2.**
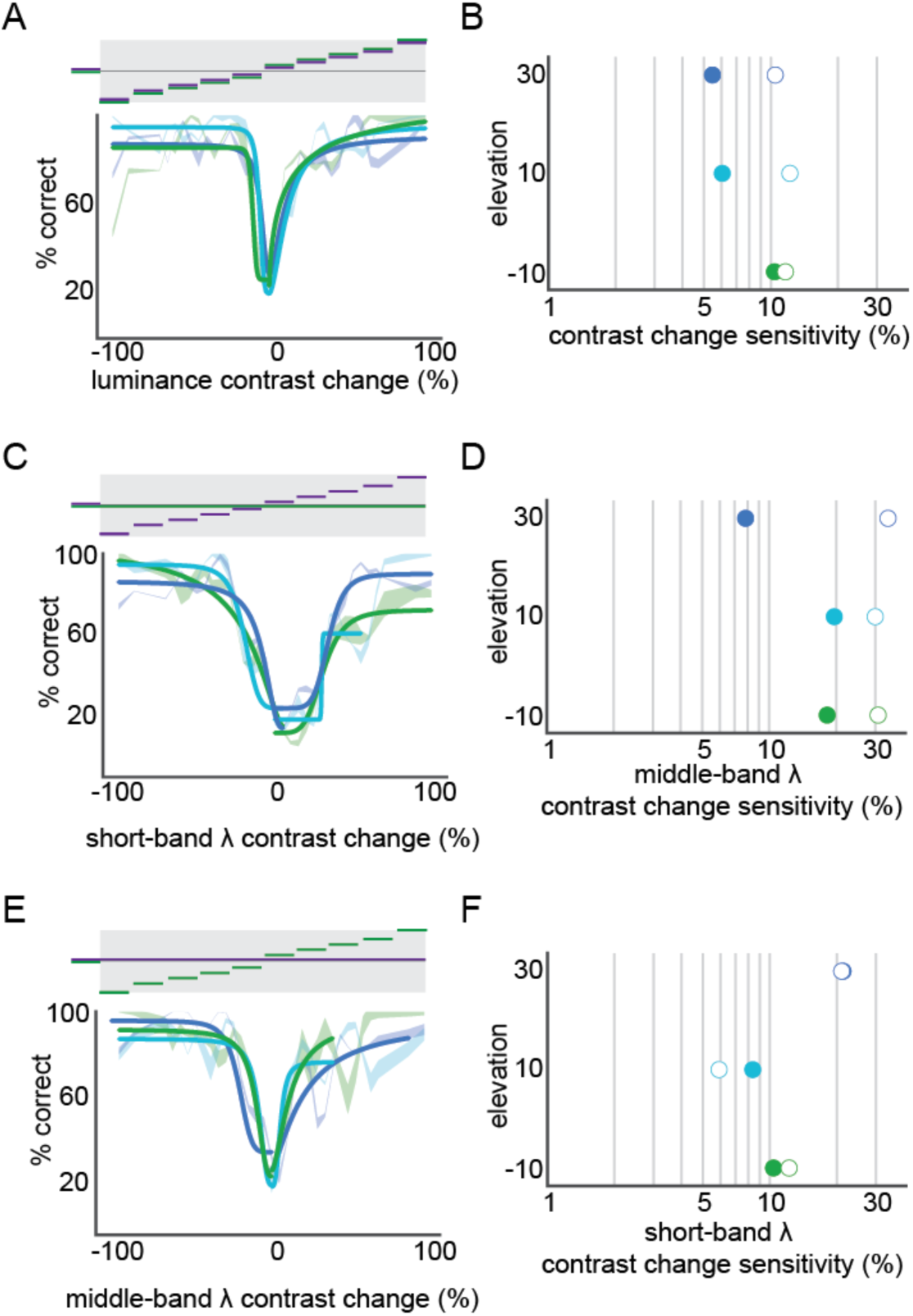
Short and middle wavelength band specific contrast sensitivity. **A**, Performance of luminance contrast change detection at three elevations (green lines: -10°, light blue lines: 10°, dark blue lines: 30°). For each elevation, a fit with hyperbolic ratio function is shown overlaid on mean performance; mean performance line thickness shows S.E.M. across mice. The stimulus is schematized above the performance, showing the corresponding relative change in each wavelength band for each condition. **B**, Contrast sensitivity at each elevation, from the fits in panel A. **C,D,** short-band specific luminance contrast performance across elevation, as in panels A,B. **E,F,** middle-band specific luminance contrast performance across elevation, as in panels A,B.

To determine the independent contributions of short and middle wavelength bands we examined change trials that contained increments or decrements of only one of the two LEDs. For the short wavelength band (i.e. UV), contrast sensitivity was non-uniform, with the highest sensitivity in the upper visual field (Fig 2C,D; 8% threshold). As with total luminance, mice were more sensitive to decrements than increments in contrast. The non-uniformity across elevation was more pronounced for short-wavelength specific sensitivity than total luminance, but was restricted to decrements. The middle-wavelength (i.e. blue/green) luminance contrast sensitivity was also non-uniform, across elevation, but with the opposite relationship as the short wavelength and total luminance. The middle-wavelength (i.e. green) luminance contrast sensitivity was nearly identical across the tested positions (Fig 2E-F). Middle-wavelength sensitivity was very similar for increments and decrements, again in contrast to total and short-wavelength luminance contrast sensitivity. In summary, we found that luminance contrast sensitivity was non-uniform, with significant opposing wavelength-band specific non-uniformities, though less than what would be predicted from the opsin expression or photoreceptor response alone.

### Determination of relative short and middle wavelength band contributions at several retinotopic locations and comparison with predicted cone weights

We next determined the relative strength of short and middle-wavelength stimulation across the visual field. Despite the measurements of the cone gradients and non-uniformity in V1 response ^23^, we were uncertain about the relative contributions of rods and cones at the tested light levels (i.e., relative contributions of the rod and middle opsin to middle band stimulation), so we first determined which combinations of wavelength band activation effectively opposed each other at each elevation.

Our approach is schematized Figure 3A: for equally weighted contributions, equal but opposite changes in the strength of the LED on the same trial should oppose each other and lead to chance change detection performance; deviations from opposing luminance changes generate luminance contrast and high detectability performance (deviations from the major axis of the ellipse). The slope of the major axis of this ellipse, hereafter called the “equiluminant line”, indicates the relative weight of each luminance band. A slope of 1 lies on the unity and equal short and middle-band weight (Figure 3A, left); slopes > 1 indicate middle band domination (Figure 3A, right) and < 1 short band domination. In setting up the visual stimulation we attempted to adjust the projector LEDs to match the strengths of short and middle band activation using published retinal sensitivities ^36^; if this were successful, we expected the axis of opposition to closely match unity.

**Figure 3.**
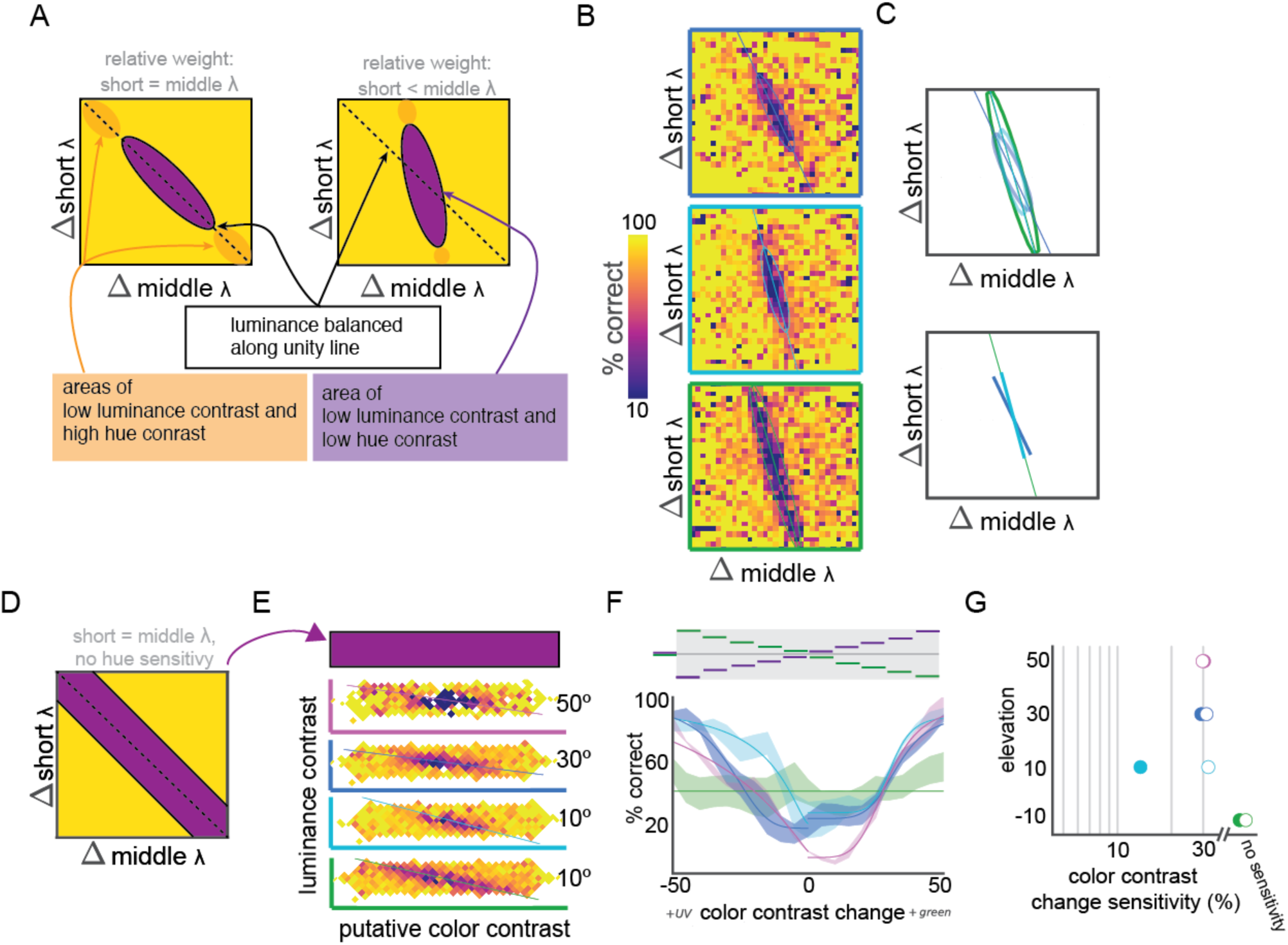
Relative short and middle-wavelength band weights and hue discrimination across elevation. **A**, Schematic representation of two possible relative short and middle band weight scenarios, with putative hue contrast sensitivity. The left scenario shows equal weight of short and middle bands, leading in balanced sensitivity and chance performance to equal and opposite changes in short and middle band contrast, along the unity line. The right scenario shows higher middle band sensitivity, resulting in a positive shift of the slope, where the larger changes in short band contrast are required to balance changes in middle band contrast. For both scenarios low luminance and hue contrast should yield to low change detection performance (purple ellipses). If hue discrimination occurs along the major axis of this purple ellipse, and should be high near the edges (orange ellipses). **B**, Observed relative weights of short and middle band weights, as in panel A., at elevations of -10° (green outline, bottom), 10° (light blue outline, middle), and 30° (dark blue outline, top). **C**, Fit of the relative weights in panel B with two-dimensional Gaussians, including a plot of the major axis of the fit. These major axes are isolated in the plot below. Color corresponds to stimulus elevation (green lines: - 10°, light blue lines: 10°, dark blue lines: 30°). **D**, schematic of sensitivity testing after attempted compensation for relative short and middle band weight. The central region should be along the line of equiluminance, so subsequent testing was focused there (arrow to panel E). **E.**, Performance of change detection at four elevations (green lines: -10°, light blue lines: 10°, dark blue lines: 30°, pink lines: 50°) after attempted balancing of short and middle weights. Because short and middle weight were not exactly balanced, performance was fit with a two-dimensional Gaussian and hue sensitivity measured along the major axis. **F**, Hue sensitivity at each elevation. Fit with hyperbolic ration function is shown overlaid on mean performance; mean performance line thickness shows S.E.M. across mice. The stimulus is schematized above the performance, showing the corresponding equal and opposite relative change in each wavelength band for each condition. **G**, Contrast sensitivity at each elevation, from fits in panel F.

We measured the axis along which opposite changes in short and middle wavelength stimulation effectively canceled each other at several elevations. The performance across pairwise combinations of changes in short and middle band luminance (Figure 3B) was fit with an ellipse (Figure 3C, top), and the major axis of this ellipse was taken to be the equiluminance line (Figure 3C, bottom). We found the mouse to be more sensitive to middle band stimulation than expected at all elevations tested, including at 30° where given the eye positioning (Figure 1 – figure supplement 1) short-opsin expression dominates. In fact, surprisingly, the mice were more sensitive to middle than short-band changes at all elevations, with the following middle/short ratios: 3.4, 3.6, and 2.25 at –10°, 10°, and 30°, respectively.

We compared our measure of the wavelength-band specific perceptual contributions to predictions from the cone expression distribution, the cone functional response, and intrinsic imaging of the mouse visual system ^15,23^. By projecting 29 the spatial profile of cones into visuotopic coordinates based on the mean eye position during our experiments (Figure 1 – figure supplement 1), we computed expected middle/short ratios for each of the elevations we tested (Figure 3 – figure supplement 1). Similar to the estimated relative opsin weights across V1 under photopic conditions ^23,37^, 2.3, 0.81, 0.81, the cone functional distribution predicts 2.3, 1.0, and 0.44 at –10°, 10°, and 30°, respectively; both predict far less middle-band sensitivity that we observed. This result suggests that rod opsin sensitivity contributes significantly to mouse perceptual sensitivity at these light levels, at least as much as 60% (Figure 3 – figure supplement 1) at higher elevations.

### Can mice discriminate hue?

We continued to ask if mice could report a change in hue independent of any luminance change. Instead of explicitly creating a device that normalized total luminance during hue changes ^38–40^, we presented sufficient combinations of wavelength-band specific luminance changes to experimentally determine when hue changes occurred independent of luminance changes. We start with the assumption that no change in luminance or hue contrast is not discriminable to the observer. All luminance and hue contrast changes that are behaviorally indistinguishable from this “no change” condition form an ellipse of non-discriminability in hue-luminance space analogous to a MacAdam ellipse of non-discriminability in color space^38^. As noted above, the axes of this ellipse that more closely matches the stimulus “luminance-balanced” line (Figure 3A, unity line) specifies this equiluminance line for the wavelength band sensitivities at that elevation; because mice are more sensitive to luminance that hue contrast, this was always the major axis of the ellipse (Figure 3A, purple ellipses). In an HSL color model, this line is equivalent to that of constant saturation and lightness but varying hue; it is also analogous to a slice through the equiluminant plane in a DKL color space^41^. Because of the shift short versus middle contributions across space, this equiluminance line changes with elevation (Fig 3C), so we attempted to create conditions of uniform luminance contrast across elevation by adjusting for the relative short and middle-band sensitivities (Figure 3D). By examining change detection performance along this experimentally defined axis of hue change, we can ask if mice can discriminate hue independent of luminance (Figure 3A, orange ellipses).

We found hue discrimination to depend on elevation. Examining the luminance-adjusted data, we found hue discrimination was negligible at -10°, but mice were capable of varying levels of hue sensitivity at all other elevations tested (Fig 3F). The performance along the equiluminant line at -10° was not well fit by a hyperbolic ratio function, and the performance in catch trials (0% contrast change) was not significantly different from any point along the line (p > .05, student’s t-test). We were able to fit the performance at each of the other elevations tested, up to 50° above the horizon. Hue contrast sensitivity was highest at for decrements in short-middle opponency at 10° elevation (13.4%); hue sensitivity was nearly identical for decrements in short-middle opponent contrast at 30 and 50° and for increments at all elevations above the horizon (29.3 – 32.1%).

## Discussion

The finding that both wavelength-specific luminance (Figure 2) and hue contrast (Figure 3E-F) is not uniform is in accordance with the distributions of both retinal and primary visual cortical ^23^ responses; however, we found that middle-band sensitivity was both higher and more uniform than expected (Figure 2). This suggests that rod sensitivity contributes significantly to perceptual sensitivity at these light levels (Figure 3 – figure supplement 1).

This finding may be important for studies of the mouse visual system that use visuotopically extended stimulation^10,25,42–44^, especially those that measure the underlying population representation of the stimuli. Because the spatial scale of luminance and contrast adaptation can be large ^45^, the adaptation to large single-band stimuli (such as those produced by LCD or other sRGB displays) in these studies may underestimate the contrast sensitivity for cells in upper visual field. This spatial scale is especially relevant because of the scale of mouse vision – 50% differences can be seen across a small number (∼5) receptive field diameters.

Our results also demonstrate that hue sensitivity depends on retinotopy, and that some retinotopic locations appear to not support hue discrimination. A goal of many large-scale data collection efforts, both completed ^18^ and underway (brain-map.org/visualcoding) as well as smaller-scale surveys ^3,13,46,47^ from retina to V1 is the classification or clustering of response properties in order to define functional channels. Because color-opponent cells, both single and double^48^, are thought to underlie such behavior, our findings indicate that mice may have at least one, likely at least two, color-opponent cell types; the presence of such functional cell types may depend strongly on retinotopy. Notably, our animals are housed in an environment with fluorescent lighting that does not provide UV-B for reflection (Figure 2 – figure supplement 1), suggesting that the behavior we observed is developmentally specified, not learned, and not lost through lack of use.

Mice, while often considered nocturnal ^49^, can be behaviorally active across a range of luminance conditions; C57BL/6 mice in particular can shift between diurnal, crepuscular, and nocturnal behavioral patterns over the course of the year ^28^. Previous studies on color signaling in the mouse have offered several hypotheses for the ethological uses of color signals. Our results are consistent with the hypothesis^15^ that non-uniform wavelength-specific sensitivity is matched to the luminance statistics of natural scenes (UV in the upper visual field, green in the lower), and high sensitivity to decrements in the short wavelengths may be particularly helpful during the shift towards UV in the spectral radiance distribution the during twilight hours ^50^. Another hypothesis, that short-middle opponency is useful for identifying mouse urine posts^17^, is inconsistent with our demonstration of a lack of hue discrimination at -10°, at least in absence of significant excursions in eye position or head movements. Our results suggest that, under the mesopic condition during which mice are often active, hue signals may be mediated by rod-cone opponency, and this may facilitate the specialization of cone opsin distributions for sampling natural luminance statistics ^15,51^.

## Methods

All procedures are approved by the Allen Institute for Brain Science Institutional Animal Care and Use Committee.

### Animals and Surgical Preparation

All animals used in this study (n=5) were C57Bl/6J male mice aged 30-300 days obtained from The Jackson Laboratories. To fix the animal’s head within the behavioral apparatus, a single surgery to permanently attach a headpost was performed. During this surgery, the animal was deeply anesthetized with 5% isoflurane and anesthesia maintained throughout the surgery with 1.5-2% continuous inhaled isoflurane. The mouse was secured in a strereotax with ear bars; hair was removed and the exposed skin sterilized with three rounds of betadine. An anterior-posterior incision was made in the skin from anterior of the eyes to posterior of the ears. The skin was removed in a tear drop shape exposing the skull. The skull was leveled and the headpost was placed using a custom stereotaxic headpost placement jig. A custom 11 mm diameter metal headpost with mounting wings was affixed to the skull using dental cement. The exposed skull inside the headpost was covered with a thin protective layer of clear dental cement and further covered with Kwikcast. The animal was allowed to recover for at least 5 days prior to the initiation of behavioral training.

After headpost implantation animals were kept on a reverse light cycle (lights OFF from 9AM to 9PM) and behavioral testing was done between 9AM and 1PM. Mice were habituated to handling gradually, through sessions of increasing duration. Mice were also habituated to the behavioral apparatus, first by allowing periods of free exploration and subsequently with head fixation sessions increasing from 10 minutes to 1 hour over the course of 1 week. Water restriction began with habituation; all mice were maintained at 85% of the original body weight for the duration of training and testing.

### Stimulus Environment and Stimuli

Ultraviolet and human-visible stimuli were provided across a range of retinotopic locations using a custom spherical stimulus enclosure ^19^ (Figure 1A, Fig 2- figure supplement 1). A custom DLP- projector designed for the mouse visual system provided independent spatiotemporal modulation of ultraviolet (peak 380nm, Figure 1B) and green (peak 532nm, Fig 1B) light. The projection system operated at 1024x768 pixel resolution and a refresh rate of 60Hz, achieving a maximum intensity of ∼3 cd/m2. Planar stimuli were spatially warped according to a custom fisheye warp for presentation on a curved screen; the fisheye warp was created through an iterative mapping protocol using the meshmapper utility (http://paulbourke.net/dome/meshmapper/) calibrated on the behavioral environment to achieve maximal accuracy.

Stimuli were presented in the right visual field and consisted of 15° diameter circles of varying color on a mean intensity background using custom written software extensions of the PsychoPy package (http://www.psychopy.org). The background intensity was 1.52 cd/m2. For some testing sessions, the color of the display was adjusted to match the mouse’s spectral sensitivity in order to create uniform and balanced sensitivity to the projector’s LED sources across the visual stimulus enclosure. To do so, a custom warp was applied that included a spatially dependent adjustment of the intensity of each LED (near-UV and ‘green’), according to the results shown in Figure 3C.

Animals were head-fixed on a freely rotating disc in the center of the spherical enclosure and allowed to run freely during the course of training and testing. A lick spout was positioned approximately 0.5 cm in front of the mouse within range of tongue extension.

In some experiments, infrared short-pass dichroic mirrors (750nm short-pass filter, <u>Edmund Optics</u>) were placed in front of each eye to allow for video tracking of the pupil. Cameras (Mako and Manta, AVT technologies) placed behind the animal were aligned to record a reflected image of the pupil; infrared illumination and a reference corneal reflection was provided via an LED positioned near the camera. Movies of the eye position during presentation of the stimuli used in the task was acquired at >=60Hz, with the eye occupying >60% of the image at 300 x 300. Data from these sessions were not included in the performance analysis to avoid any potential artifact caused by the infrared dichroic.

### Behavioral Task

Animals were first shaped to associate changes in luminance with a reward. After each change in luminance a water reward was automatically delivered, regardless of mouse licking behavior. During these sessions, the reward was constant at 10µL. Incorrect licks were punished by resetting the trial, such that the mouse had to wait longer for the next change. This “shaping” phase lasted a minimum of two days, but for most mice extended to several weeks. For some animals (2/5), subsequent epochs of this automatic reward shaping served as task reinforcement when performance in testing blocks dropped.

During each testing session a circle was presented at a single visuotopic location and remained at this location for the duration of the session. At non-regular intervals, again selected from an exponential distribution, the color and/or luminance of the stimulus was changed, and the mouse had to report detection of change by licking the reward spout within 1 second of the change in order to receive reward (Figure 1C). Licks were detected through a capacitive sensor connected to the reward spout. No water was present on the reward spout before the first lick; if the animal correctly detected a change, a water reward (3-10µL, depending on animal and stage of training) was delivered through this spout (Figure 1D). Sessions were 50-60 minutes and typically included ∼300 trials.

Mice were first trained to associate changes in a 15° stimulus at 100% luminance contrast with a reward. In these sessions (total of 3 to 25 sessions), the contrast of a stimulus (10° elevation, relative to the horizon) changed at exponentially distributed intervals from 50% positive relative to the background to 50% negative (from white to black), or vice versa (black to white). If a lick occurred within 1 second of an actual stimulus change (Figure 1B), a reward was delivered to the spout and liquid reward was consumed subsequent licks (Figure1C). If a lick occurred outside of this window the trial was aborted, extending the time the mouse must wait and effectively creating a ‘time-out’ period. Mice advanced from this protocol after performance exceeded 75% for consecutive sessions.

In subsequent testing sessions, the intensity of the ultraviolet and green intensities were varied independently on each trial. Each trial contained a change in LED intensity for a 15° test circle on a mean luminance background at one of four elevations: -10°, 10°, 30°, and in some cases 50°. The first 8-20 trials of each session were 100% contrast changes, as described for the training blocks, with rewards automatically delivered. The number of these daily “free” rewards was reduced to 8 for as long as the mouse received >1.0mL of reward during training or performed well enough to reach satiety and disengage from the task. We attempted to correct for sessions with poor performance by increasing these “free” rewards on subsequent days before gradually reducing them again. To control for motivation in the results, we calculated a running average of the reward rate and selected trials where this reward rate remained above 4 rewards per minute; only these engaged trials (44%, 56112/127659) were used for analysis.

### Analysis

All analyses were done using Python and common scientific packages (numpy, scipy, matplotllib, and pandas). Code is publicly available from github.com/danieljdenman/mouse_chromatic and includes a Jupyter notebook that contains code for generation of our figures from the data. Data from each training session was saved and combined into a common data structure that was used for all analysis. Individual sessions were analyzed to drive adjustments in the training parameters such as the number of automatic “free” rewards. Following data collection for all animals, all sessions were loaded into a single object for analysis. This data structure can be recreated from the NWB^52^ files made available from <github.com/danieljdenman/mouse_chromatic.>.

To quantify performance, from each trial the following parameters were extracted: change times (the time of stimulus change), lick times (the time of each lick, as detected through the capacitive sensor connected to the reward spout), and the stimulus conditions. A trial was scored “correct” if the first lick after a change time occurred with one second, and if there was actually change in intensity of either green or ultraviolet at that change time.

For each mouse, the percent correct was computed for each pair of LED state transitions, i.e., each pairwise combination of change in short-band luminance and change in middle-band luminance (e.g., Fig 3C). For each mouse, performance was ignored if 3 trials were not presented for those conditions. For fitting, missing data were replaced via a nearest neighbors approach, with the mean of the surrounding data. Our sampling strategies focused on the areas of changing performance, ensuring that cases of missing data were limited to the areas where performance had saturated at or near the lapse rate. Psychophysical curves for wavelength band-specific and hue sensitivity were taken from the appropriate slices of this color space. Sensitivity was taken from the c50 parameter of fit a hyperbolic ratio fit^53^.

A total of 5 mice entered training on the task; one mouse failed to reach consecutive sessions of 75% performance during the initial high luminance contrast change detection phase, and so did not continue to testing in the hue contrast discrimination phase. We did not use any statistical methods to determine mouse or trial sample size prior to the study, determining based on stability and consistency of results when sufficient samples had been collected. Statistical tests were student’s t-test unless otherwise specified.

Eyetracking analysis was done via a semi-automated algorithm; full details are available from <http://help.brain-map.org/display/observatory/Documentation>. Briefly, the algorithm fits an ellipse to the pupil or corneal reflection (CR) area, respectively. A seed point is identified by convolution with a black square (for the pupil) and white square (for the corneal reflection). An ellipse was fit to candidate boundary points identified using ray tracing using a random sample consensus algorithm. The fit parameters were first reported in coordinate centered on the mouse eye, and subsequently converted to visual degrees by projection based on the position of the dichroic mirror and the relative position of pupil and corneal reflection. Coordinates for eye position were extracted independently for each frame of the eye position movie.

## Acknowledgements

We would like to thank Saskia de Vries and Brian Long for useful discussions in the preparation of this manuscript. We would also like to thank the Neurosurgery and Behavior team, Animal Care team, and Naveen Oullette for assistance in animal surgery, care, and handling. We wish to thank the Allen Institute founders, Paul G Allen and Jody Allen, for their vision, encouragement and support.

## Contributions

Conceptualization, DJD and RCR; Methodology, DO, SO; Investigation, DJD, JL, and SC; Software, DJD, DW, DO, and MB; Writing – Original Draft, DJD and RCR; Writing – Review and Editing, DJD, JL, DO, SC, DW, MB, SO, and RCR.

---

**Figure 1 – figure supplement 1.**
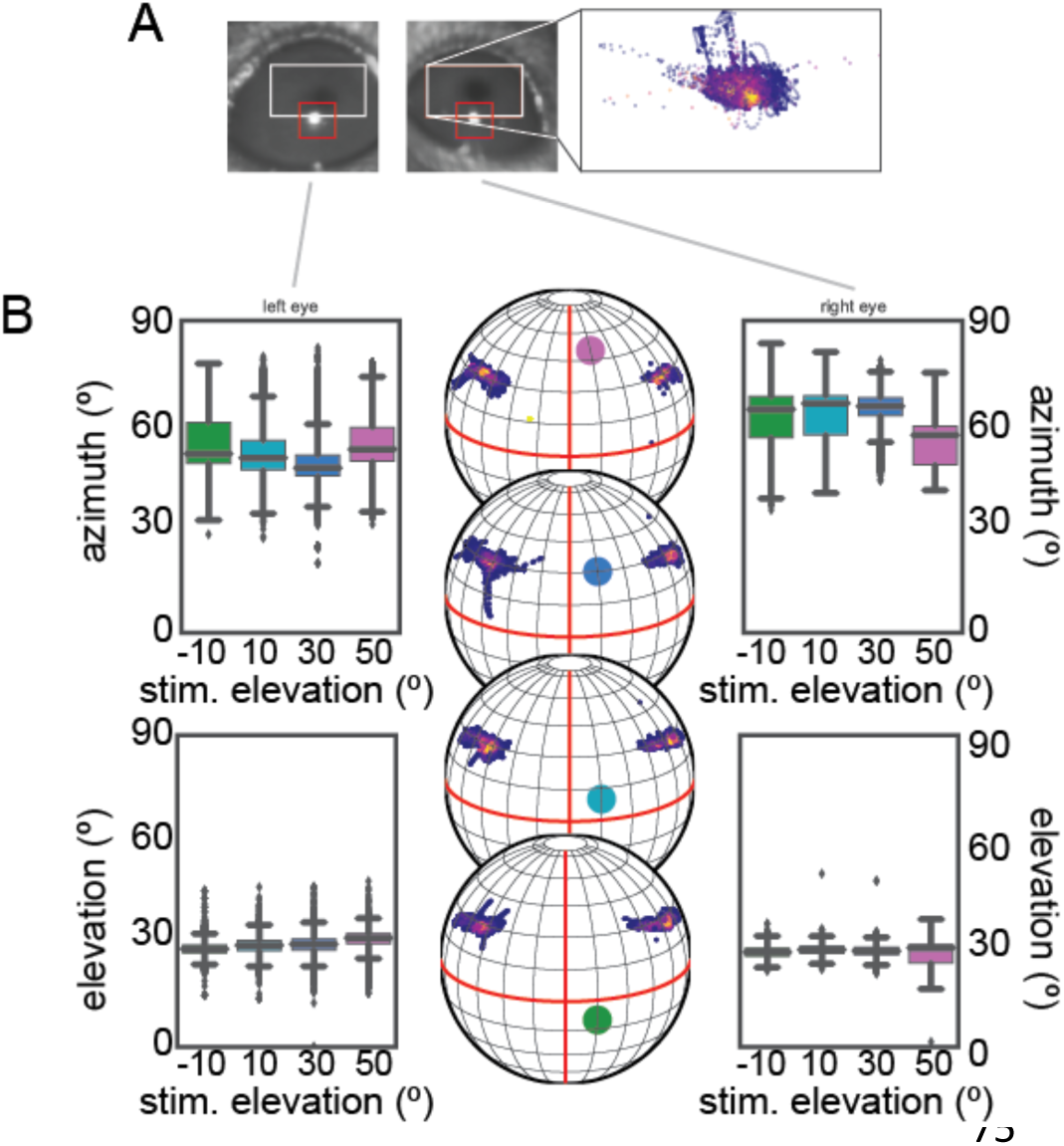
Eye position during performance of the change detection task at four elevations. **A,** Example images from the pupil tracking of the left eye (left) and right eye (right). For both eyes, user specified windows specifying the location of the pupil and corneal reflection are overlaid on the images; for the right eye, the center position of an ellipse fit to the pupil is shown within this box, with the color of each point representing the density of overlapping points. **B,** Eye position across the four elevation conditions. The elevation and azimuth are not statistically different between any pair of conditions (p > 0.05, t-test).

**Figure 2 – figure supplement 1.**
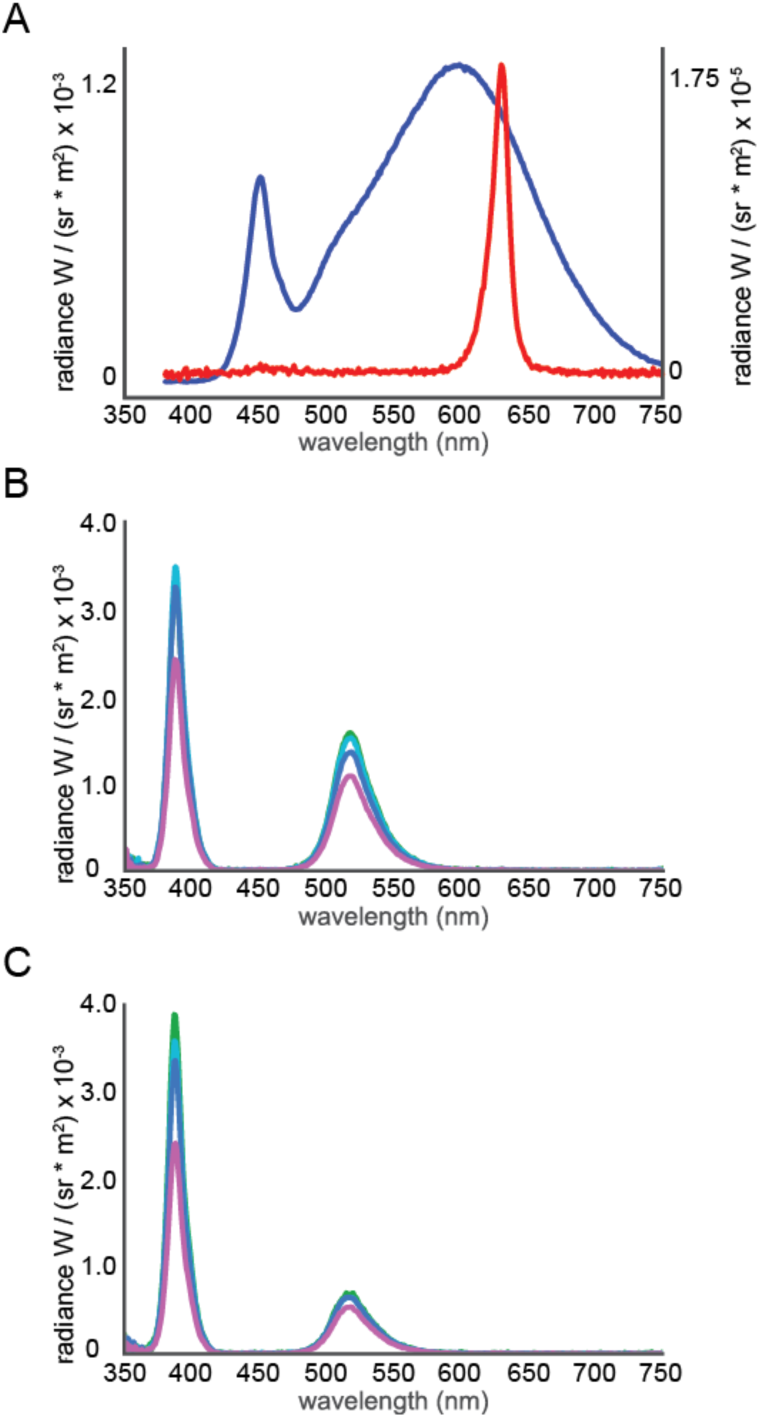
Spectral radiance of various illumination sources from 350 – 750nm. **A,** The spectral radiance of the lighting in the Allen Institute animal housing facility, including the behavioral testing suite during lights on (blue line, left axes) and lights off (red line, right axes) conditions. Note the lack of any irradiance below 400nm. **B,** The spectral radiance off of the stimulation dome at each elevation for equal short and middle LED drive **C**, The spectral radiance off of the stimulation dome at each elevation after adjustment for wavelength-band Specific non-uniformity.

**Figure 3 – figure supplement 1.**
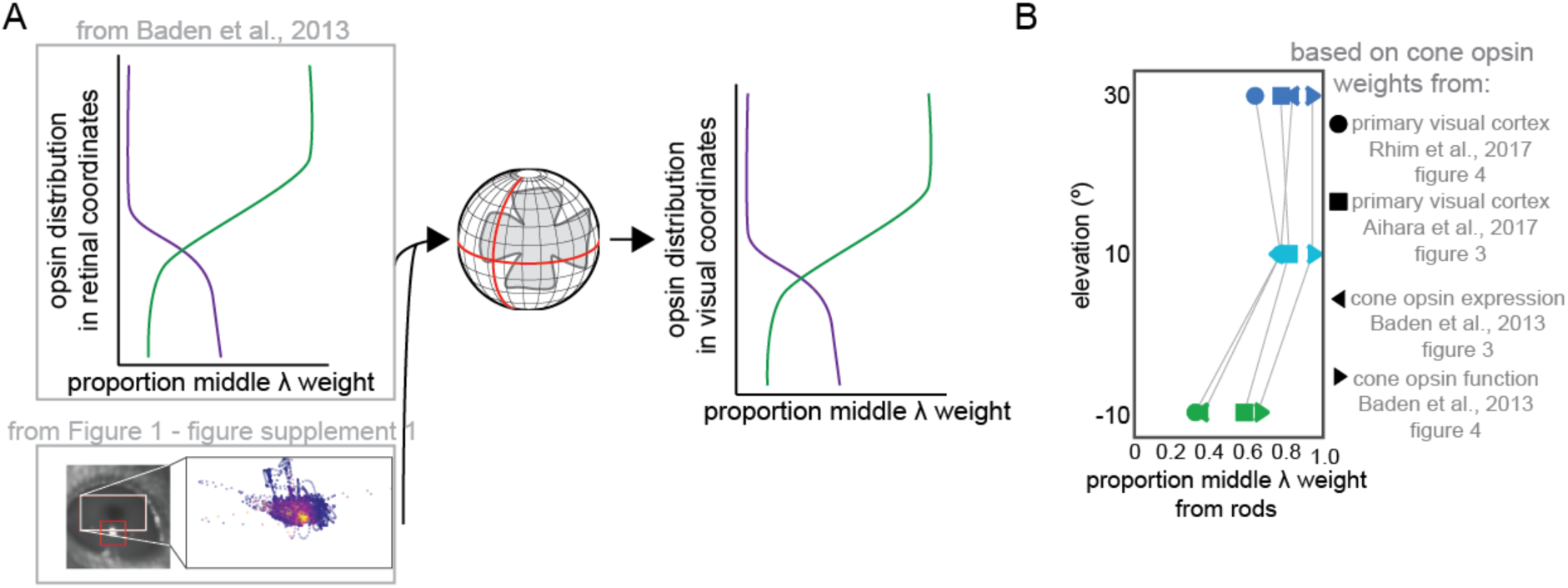
Estimating short and middle cone weights in the coordinates of our behavioral apparatus from retinal expression and functional data. **A**, To generate predictions of the relative weights of each wavelength band at each elevation tested in our paradigm, we projected retinal spatial distributions of both opsin expression and functional responses, both from Baden et al., 2013, into spherical coordinates according to the eye position measured in Figure 2 – figure supplement 1. **B,** normalized difference between the predicted (cone opsin expression, cone opsin functional response, V1 response) and observed behavioral weights, for -10°, 10, and 30°. The difference between the behavior middle/short ratio at each elevation and the middle/short ratios at those elevation predicted by several measures of cone opsin weight in the literature was divided by the observed ratio to estimate the proportion of the middle band weight provided by rods under our behavioral luminance conditions

